# Suramin inhibits SARS-CoV-2 infection in cell culture by interfering with early steps of the replication cycle

**DOI:** 10.1101/2020.05.06.081968

**Authors:** Clarisse Salgado Benvindo da Silva, Melissa Thaler, Ali Tas, Natacha S. Ogando, Peter J. Bredenbeek, Dennis K. Ninaber, Ying Wang, Pieter S. Hiemstra, Eric J. Snijder, Martijn J. van Hemert

## Abstract

The SARS-CoV-2 pandemic that originated from Wuhan, China, in December 2019 has impacted public health, society and economy and the daily lives of billions of people in an unprecedented manner. There are currently no specific registered antiviral drugs to treat or prevent SARS-CoV-2 infections. Therefore, drug repurposing would be the fastest route to provide at least a temporary solution while better, more specific drugs are being developed. Here we demonstrate that the antiparasitic drug suramin inhibits SARS-CoV-2 replication, protecting Vero E6 cells with an EC_50_ of ∼20 µM, which is well below the maximum attainable level in human serum. Suramin also decreased the viral load by 2-3 logs when Vero E6 cells or cells of a human lung epithelial cell line (Calu-3) were treated. Time of addition and plaque reduction assays showed that suramin acts on early steps of the replication cycle, possibly preventing entry of the virus. In a primary human airway epithelial cell culture model, suramin also inhibited the progression of infection. The results of our preclinical study warrant further investigation and suggest it is worth evaluating whether suramin provides any benefit for COVID-19 patients, which obviously requires well-designed, properly controlled randomized clinical trials.

## Introduction

In December 2019, local health authorities reported an increasing number of pneumonia cases, rapidly spreading across the city of Wuhan, Hubei province, in China (1). Further analysis showed that the causative agent of this disease was SARS-coronavirus-2 (SARS-CoV-2), which is a member of the betacoronavirus genus within the coronavirus family and shares roughly 80% of genetic identity with SARS-CoV (2, 3). Since then, SARS-CoV-2 has spread to 113 countries, leading to a coronavirus pandemic of unprecedented magnitude, with more than 3.5 million confirmed cases globally and more than 240,000 casualties reported by WHO on May 5^th^, 2020 (4).

Coronaviruses are enveloped viruses, that possess extraordinarily large (26 to 32 kb) positive-strand RNA genomes (5). SARS-CoV-2 infection often causes only mild disease, but can also lead to clinical manifestations such as high fever, cough, dyspnea, myalgia and headache. Although the majority of cases may be asymptomatic or present mild symptoms with good recovery, some patients develop more severe outcomes, such as severe pneumonia, respiratory failure, multiple organ failure or death (6).

Due to the urgency of the situation, the lack of approved specific antiviral therapy against coronaviruses and the time it takes to develop the latter through regular preclinical and clinical research, there is great interest in repurposing already approved drugs. This would be a fast track to apply candidate therapeutic agents as antivirals to combat SARS-CoV-2 infection, which can be used to fight the virus while better and more specific antivirals are being developed.

Drugs like ribavirin, remdesivir, favipiravir and the anti-malarial therapeutic chloroquine showed promise in cell culture and some also appeared to show (modest) effects in early trials in humans, which were not always conducted with the most optimal design (7). However, except for remdesivir (8) more recent (and more appropriately conducted) clinical trials suggest that none of these drugs provide substantial benefit in patients and that they should be used with caution due to their potential side-effects. Therefore, it appears that options to inhibit SARS-CoV-2 infection are limited and mainly supportive care and treatments that target the immune system and inflammatory responses can be provided to patients. This stresses the urgency of evaluating additional approved drugs as candidates for use as antiviral therapy against this pathogen.

We now provide evidence showing that suramin can be considered as drug candidate that deserves further assessment, as we found the compound to exhibit antiviral activity against SARS-CoV-2 in relevant cell culture models at concentrations that can be easily reached in human serum. Suramin is an anti-parasitic drug that is used to treat sleeping sickness caused by trypanosomes. It is a symmetrical polysulfonated compound that was synthesized for the first time around 1916 (9). Later we and many others have shown that suramin also has broad-spectrum antiviral effects, as it inhibits HIV (10), hepatitis C virus (11), herpes simplex type-1 virus (12), Zika virus (13), dengue virus (14), chikungunya virus (15), and others.

In the present study, we show that suramin also exhibits antiviral activity against SARS-CoV-2 in cell culture, most likely by inhibiting viral entry. The compound had an EC_50_ of 20 µM in Vero E6 cells and showed a more than 2 log viral load reduction when infected human Calu-3 airway epithelial cells were treated. Finally, suramin reduced SARS-CoV-2 progression of infection in well-differentiated primary human airway epithelial cells cultured at the physiological air-liquid interface. It is important to stress that these results should not be directly translated to efficacy against SARS-CoV-2 in humans and guarantee no benefit to the patient yet. However, our results make suramin an interesting candidate to further evaluate in in-depth pre-clinical studies (e.g. into formulation, mode of administration, pharmacokinetics and in other *ex vivo* models) and suggest suramin could be explored in carefully performed and properly controlled clinical trials for the treatment of COVID-19 patients.

## Material and Methods

### Cell lines, virus and compound

Vero E6 cells were maintained in Dulbecco’s modified Eagle’s medium (DMEM; Lonza), supplemented with 8% fetal calf serum (FCS; Bodinco), 2 mM L-glutamine, 100 IU/ml of penicillin and 100 µg/ml of streptomycin (Sigma-Aldrich). The human lung epithelial cell line Calu-3 2B4 (referred to as Calu-3 cells) was maintained as described (16). Primary human airway epithelial (HAE) cell cultures were established at the Leiden University Medical Center (LUMC; Department of Pulmonology) and their culture and infection are described below. All cell cultures were maintained at 37°C in an atmosphere of 5% CO_2_ and 95%–99% humidity. Infections were performed in Eagle’s minimal essential medium (EMEM; Lonza) with 25 mM HEPES (Lonza), further supplemented with 2% FCS, L-glutamine (Sigma-Aldrich), and antibiotics.

The clinical isolate SARS-CoV-2/Leiden-002 was isolated from a nasopharyngeal sample at LUMC and its sequence and characterization will be described elsewhere (manuscript in preparation). SARS-CoV-2/Leiden-002 was passaged twice in Vero E6 cells and virus titers were determined by plaque assay as described before (17). Working stocks yielded titers of 5 × 10^6^ plaque forming units (PFU)/ml. All experiments with infectious SARS-CoV-2 were performed in a biosafety level 3 facility at the LUMC. Suramin was purchased from Sigma-Aldrich and was dissolved in milliQ and stored at −20°C. Addition of compound to Vero E6 and Calu-3 cells was done in infection medium and in PBS for HAE cultures.

### Human airway epithelial cell cultures (HAE)

HAE cell cultures were cultured as previously described (18). Briefly, primary human bronchial epithelial cells were isolated from tumour-free resected bronchial tissue from patients undergoing resection surgery for lung cancer at the LUMC. Use of such lung tissue that became available for research within the framework of patient care was in line with the “Human Tissue and Medical Research Code of conduct for responsible use” (2011) (www.federa.org), which describes the opt-out system for coded anonymous further use of such tissue. To achieve mucociliary differentiation, PBEC were cultured at the air-liquid interface (ALI) for 21 days as previously described (18, 19). In brief, expanded HAE cells from 3 donors at passage 2 were combined (3 × 10^4^ cells per donor) and were seeded on 12-well transwell membranes (Corning Costar), which were coated with a mixture of BSA, collagen type 1, and fibronectin. In addition, cells from two individual donors were seeded on separate sets of transwell membranes. BEpiCM-b:DMEM (B/D)-medium (1:1) was used as described (supplemented with 12.5mM HEPES, bronchial epithelial cell growth supplement, antibiotics, 1 nM EC23 (retinoic acid receptor agonist), and 2 mM glutaMAX). After confluence was reached, cells were cultured at the ALI in complete medium with 50 nM EC23 for 21 days. The mucociliary differentiated cultures were characterized by a high trans-epithelial electrical resistance (TEER>500 Ω·cm2), visible cilia beating and mucus production. Before infection, cells were incubated overnight in the BEpiCM-b/DMEM 1:1 medium mixture from which EGF, BPE, BSA and hydrocortisone were omitted and that did contain antibiotics (starvation medium).

### RNA isolation and quantitative RT-PCR (RT-qPCR)

RNA was isolated from cell culture supernatants and cell lysates using the TriPure Isolation Reagent (Sigma-Aldrich). Equine arteritis virus (EAV) in AVL lysis buffer (Qiagen) was spiked into the reagent as internal control for extracellular RNA samples. The cellular household gene PGK-1 served as control for intracellular RNA. Primers and probes for EAV and PGK1 and the normalization procedure were described before (20). Viral RNA was quantified by RT-qPCR using the TaqMan(tm) Fast Virus 1-Step Master Mix (Thermo Fisher Scientific). Primers and probes were based on (21) but with modifications resulting in the following primer and probe sequences: SARS-CoV-2 N-Gene Fwd-CACATTGGCACCCGCAATC, Rev-GAGGAACGAGAAGAGGCTTG and Probe YakYel-ACTTCCTCAAGGAACAACATTGCCA-BHQ1; RdRp-Gene Fwd-GTGARATGGTCATGTGTGGCGG, Rev-CARATGTTAAASACACTATTAGCATA and Probe FAM-CCAGGTGGAACMTCATCMGGWGATGC-BHQ1. A standard curve of 10-fold serial dilutions of a T7 RNA polymerase-generated *in vitro* transcript containing the RT-qPCR target sequences was used for absolute quantification. A RT-qPCR program of 5 min at 50 °C and 20 s at 95 °C, followed by 45 cycles of 5 s at 95 °C and 30 s at 60 °C, was performed on a CFX384 Touch(tm) Real-Time PCR Detection System (Bio-Rad).

### Cytopathic effect (CPE) reduction assay

CPE reduction assays were performed as described (22). Briefly, Vero E6 cells were seeded in 96-well cell culture plates at a density of 10^4^ cells per well. Cells were incubated with 1.7-fold serial dilutions of suramin starting from a concentration of 120 µM. Subsequently, cells were either mock-infected (analysis of cytotoxicity of the compound) or were infected with 300 PFU of virus per well (MOI of 0.015) in a total volume of 150 µl of medium. Cell viability was assessed three days post-infection by MTS assay using the CellTiter 96^®^ Aqueous Non-Radioactive Cell Proliferation kit (Promega) and absorption was measured at 495 nm with an EnVision Multilabel Plate Reader (PerkinElmer). The 50% effective concentration (EC_50_), required to inhibit virus-induced cell death by 50%, and the 50% cytotoxic concentration (CC_50_), that reduces the viability of uninfected cells to 50% of that of untreated control cells, were determined using non-linear regression with GraphPad Prism v8.0.

### Viral load reduction assays

Cells were seeded in 96-well cell culture plates at a density of 10^4^ (Vero E6) or 6 × 10^4^ (Calu-3) cells per well in 100 µl culture medium. As control to determine the amount of residual virus after removal of the inoculum and washing, cells in some wells were killed with 70% Ethanol (followed by washing with PBS). Vero E6 and Calu-3 cells were incubated with 2-fold serial dilutions of a starting concentration of 200 µM of suramin and subsequently infected with 2 × 10^4^ PFU of SARS-CoV-2 (MOI of 1 on Vero E6 cells). For analysis of viral RNA, supernatant was harvested from Vero E6 cells at 16 h.p.i and from Calu-3 cells at 21 h.p.i. Intracellular RNA was collected by lysing the cells in 150 µl Tripure reagent. Analysis of viral progeny in supernatant from Calu-3 cells was performed by plaque assay on Vero E6 cells (17). Potential cytotoxicity of the compound was tested in parallel on uninfected cells using the MTS assay (Promega) as described for the CPE reduction assay.

### Entry inhibition plaque reduction assay

A day before infection Vero E6 cells were seeded in 6-well cell culture plates at a density of 3.5 × 10^5^ cells per well in 2 ml medium. 10^−2^ to 10^−5^-fold serial dilutions of a SARS-COV-2 stock were prepared in medium containing 100, 50, 25, 12.5, 6.25 or 0 µM suramin. These were used as inoculum to infect the Vero E6 cells in 6-well clusters. After 1 h at 37°C, the inoculum was removed and cells were incubated in Avicel-containing overlay medium without suramin for 3 days, after which they were fixed with 3.7% formaldehyde, stained with crystal violet and plaques were counted (17).

### Time of addition assay

Vero E6 cells were seeded in 24-well clusters at a density of 6 × 10^4^ cells per well. The next day cells were treated with 100 µM suramin during the time intervals indicated in Fig. 3 and they were infected at an MOI of 1. Supernatant was harvested at 10 h.p.i. for quantification of viral RNA by RT-qPCR.

### Infection and suramin treatment of HAE cells

The apical sides of HAE cell cultures were washed 3 times with 200 µl PBS for 10 min at 37°C on the day before infection to remove excess mucus. Washing was repeated once before cells were infected on the apical side with 3 × 10^4^ PFU SARS-CoV-2 (estimated MOI of 0.1) in 200 µl of PBS. The apical side was treated with 100 µM suramin in 50 µl of PBS at 12 and 24 h.p.i (after first collecting a 200 µl PBS wash to determine viral load). Control wells were treated with 50 µl of PBS. The experiment was done in triplicate, with one insert (transwell) containing a mix of cells from 3 donors and two ‘single donor’ inserts seeded with cells from two different donors. Supernatants were collected from infected PBS-treated cells and infected suramin-treated cells at 12, 24 and 48 h.p.i, by incubating the apical side with 200 µl PBS for 10 min at 37°C and collecting it. This supernatant was used for quantification of viral RNA by RT-qPCR and viral load (infectivity) by plaque assay on Vero E6 cells. At each timepoint cell lysates were collected from inserts by adding 750 µl Tripure reagent. Assessment of potential cytotoxicity of the 48h suramin treatment, compared to PBS treatment, was done with uninfected cells by MTS assay (Promega) and LDH Assay (CytoTox 96^®^ Non-Radioactive Cytotoxicity Assay, Promega) according to the manufacturer’s instructions.

## Results

### Suramin inhibits SARS-CoV-2 replication in Vero E6 cells

To determine if suramin could protect cells from SARS-CoV-2 infection and to evaluate its toxicity, Vero E6 cells were infected with SARS-CoV-2 and treated with serial dilutions of suramin in a CPE reduction assay. Suramin protected infected cells from SARS-CoV-2-induced cell death in a dose-dependent manner, with an EC_50_ of 20 ± 2,7 µM (Fig. 1A). In parallel, non-infected cells were treated with the same concentrations of suramin in order to assess the compound’s toxicity. No toxicity was observed over the range of concentrations that was used in these antiviral assays. Only at 5 mM cell viability dropped to 67%, resulting in a CC_50_ > 5 mM (15). Therefore, suramin inhibits SARS-CoV-2 with a selectivity index (SI) higher than 250.

**Figure 1.**
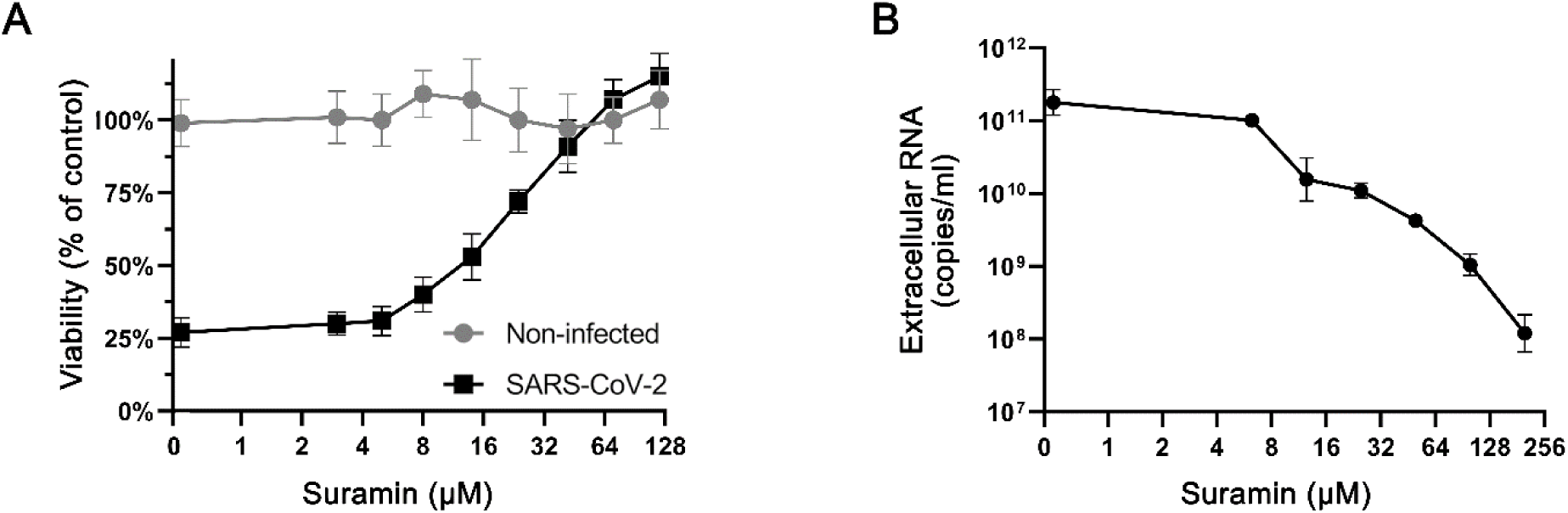
Suramin inhibits SARS-CoV-2 replication in Vero E6 cells. (A) CPE reduction assay. Vero E6 cells were infected with SARS-CoV-2 at an MOI of 0.015 and were treated with 1.7-fold serial dilutions of suramin. Viability was measured by MTS assay at 3 days post infection. The viability of non-infected suramin-treated cells was measured in parallel to assess toxicity (3 independent experiments performed in quadruplicate). (B) Viral load reduction assay. Vero E6 cells were infected at an MOI of 1, followed by treatment with different concentrations of suramin. After 16 hours, supernatants were harvested and the viral load was determined by quantification of extracellular SARS-CoV-2 RNA by an internally controlled multiplex RT-qPCR (n=3).

To more directly measure the inhibition of viral replication by suramin, viral load reduction assays were performed. Vero E6 cells were infected with SARS-CoV-2 at an MOI of 1 and they were treated with increasing concentrations of suramin. At 16 h.p.i., supernatant was harvested to determine the viral load by quantifying the levels of extracellular viral RNA by RT-qPCR (Fig 1B). The supernatant of untreated infected cells contained 10^11^ copies/ml of viral RNA. RT-qPCR revealed that the RNA levels decreased upon suramin treatment in a dose-dependent manner, showing a 3-log reduction at the highest concentration tested (200 µM) (Fig. 1B). Together, these results indicated that suramin protects Vero E6 cells from the SARS-CoV-2-induced cytopathic effect and that it reduces the viral load in these cells.

### Suramin reduces the viral RNA and infectious virus load in cultured human lung epithelial cells

To assess the antiviral effect of suramin in a more relevant model, human lung epithelial cells (Calu-3) were infected with 2 × 10^4^ PFU of SARS-CoV-2 in the presence of 0-200 µM suramin for 1h. After removal of the inoculum and washing of the cells, incubation was continued in medium with suramin (0-200 µM) for 20 hours. At 21 h.p.i., RNA was isolated from cells and supernatant and the viral titer in the supernatant was determined by plaque assay. We observed a strong dose-dependent reduction in intracellular (Fig. 2A) and extracellular (Fig. 2B) viral RNA levels in suramin-treated samples. At 200 µM the extracellular viral RNA levels showed a 3-log reduction, while intracellular viral RNA levels decreased by 2-log. Figures 2A and 2B show the results of RT-qPCR reactions targeting the RNA-dependent RNA polymerase coding region, but similar reductions in copy numbers were observed with RT-qPCR reactions targeting the SARS-CoV-2 N protein gene (also detects subgenomic RNA), although in that case absolute copy numbers -as expected-were higher than for genomic RNA (data not shown). Plaque assays confirmed that treatment with 200 µM suramin led to an almost 3-log drop in infectious progeny titers from infected-Calu-3 cells (Fig. 2C).

**Figure 2.**
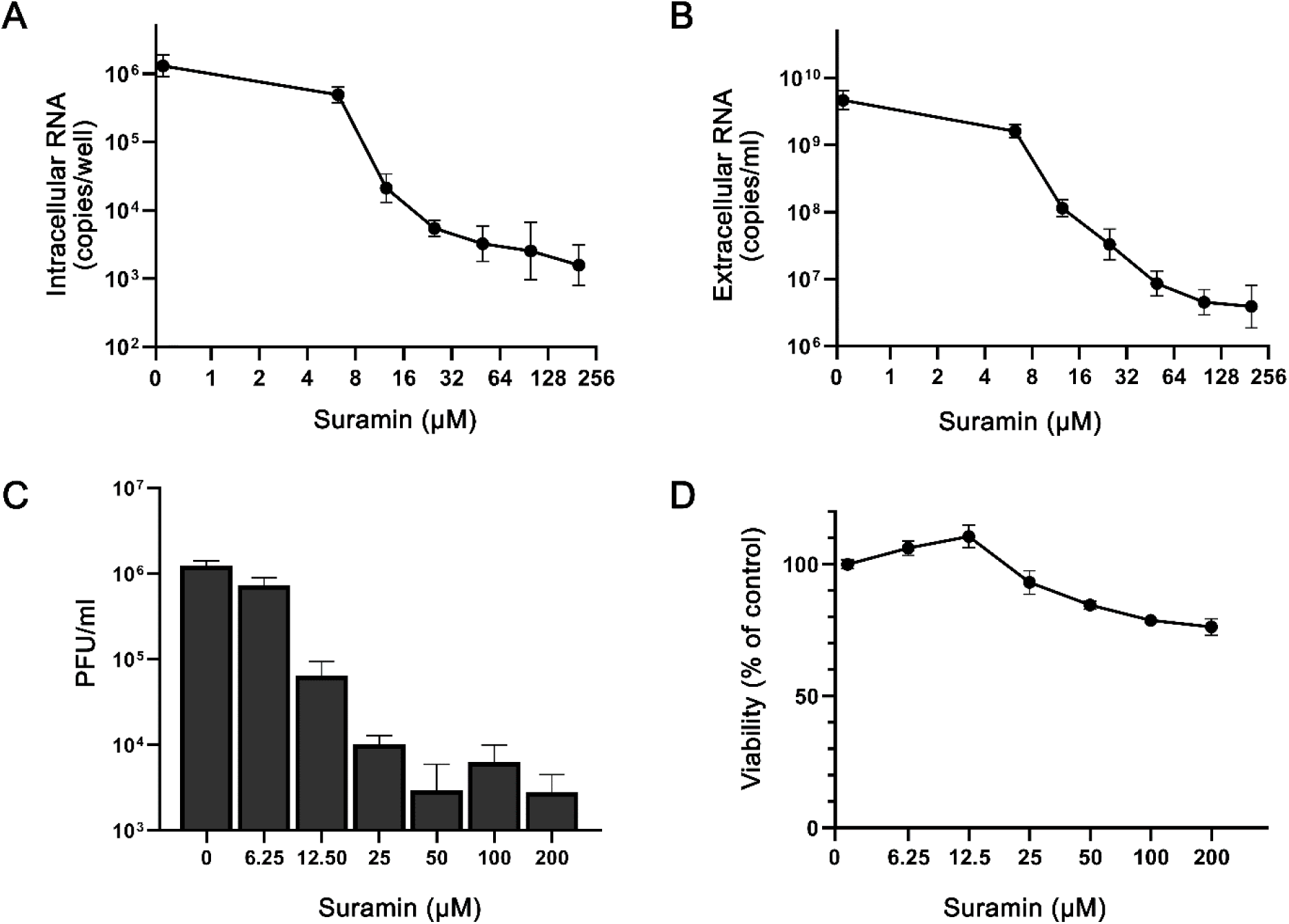
Suramin decreases levels of intra- and extracellular viral RNA and infectious progeny in infected Calu-3 cells. Calu-3 cells were infected with SARS-CoV-2, followed by treatment with 0-200 µM suramin. (A) Intracellular viral RNA copy numbers at 21 h.p.i., determined by internally controlled multiplex RT-qPCR targeting the SARS-CoV-2 RdRp coding region and using the housekeeping gene PGK1 for normalization. (B) Extracellular viral RNA levels at 21 h.p.i., quantified by RT-qPCR. (C) Viral load in the supernatant at 21 h.p.i. as determined by plaque assay on Vero E6 cells. (D) Viability of uninfected Calu-3 cells treated with various concentrations of suramin measured by MTS assay in parallel to the infection (n=3).

**Figure 3.**
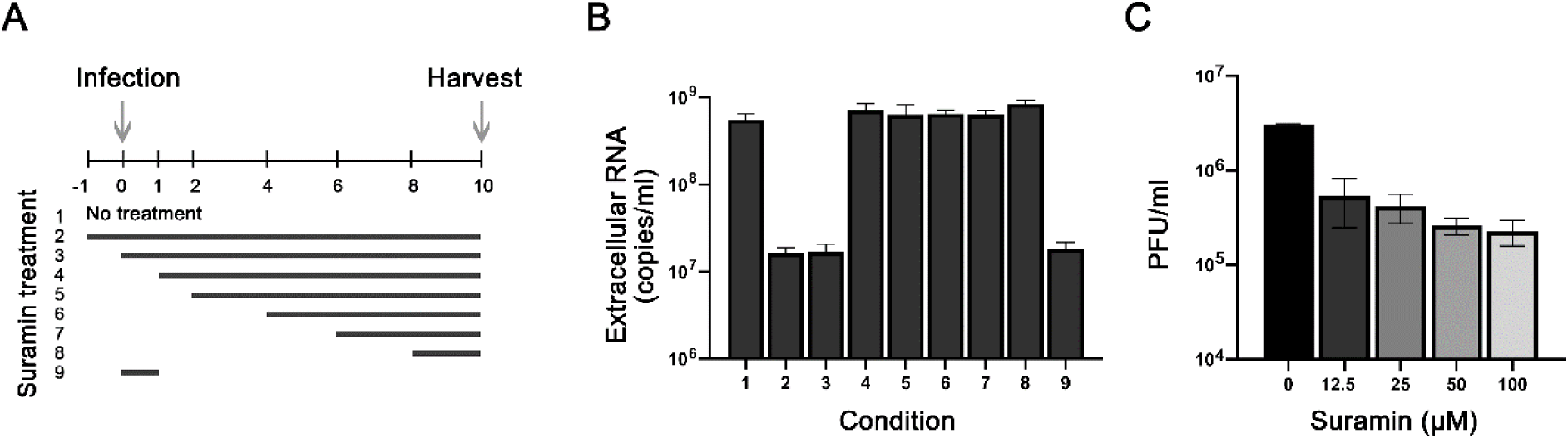
Suramin inhibits the early steps of SARS-CoV-2 replication. (A) Schematic representation of the time-of-addition experiment and the different treatment intervals. (B) At 10 h.p.i. supernatant was harvested and extracellular viral RNA levels were determined by RT-qPCR. (C) Vero E6 cells were infected with SARS-CoV-2 in the presence of various concentrations of suramin. Suramin was only present during the one hour of infection and after 1 hour, cells were incubated in overlay medium without suramin. After three days, cells were fixed, stained and plaques were counted. (n=3).

Cytotoxicity assays performed in parallel in non-infected Calu-3 cells showed that suramin was slightly more toxic to these cells than to Vero E6 cells, although cell viability remained above 80% even at the highest dose tested (Fig. 2D). Together these results suggest that suramin is a potent SARS-CoV-2 inhibitor with high selectivity, also in human lung cells.

### Suramin acts on the early steps of viral replication

To determine which step of viral replication is affected by suramin, we performed a time-of-addition assay. Cells were infected with SARS-CoV-2 (MOI 1) and treated with 100 µM of suramin over different time intervals, as schematically depicted in Figure 3A. Treatment was initiated 1 hour before infection or at 0, 1, 2, 4, 6 or 8 h.p.i., and suramin remained present until 10 h.p.i., when supernatants were harvested to determine viral load by RT-qPCR targeting the RdRp coding region. In one sample suramin was only present for 60 min during the time of infection. After 1 hour, virus inoculum was removed and cells were washed three times with PBS, followed by incubation in medium with or without suramin. At 10 h.p.i., supernatant was collected to evaluate the levels of viral RNA (Fig. 3B). When suramin treatment was initiated 1 hour before (−1h) or at the time of infection (0h) a 2-log reduction in viral RNA levels was observed. Treatments that started later than 1 hour post infection did not inhibit viral replication, as viral RNA levels similar to the non-treated control were observed. Treatment only during the infection (0-1h) resulted in the same 2-log reduction in viral RNA load as the 0-10h treatment, indicating that suramin inhibits an early step of the replication cycle, likely viral entry.

To confirm suramin’s inhibitory effect on entry, we performed a plaque reduction assay, by infecting Vero E6 cells with serial dilutions of SARS-CoV-2 in the presence of increasing concentrations of suramin, which was only present during the one hour of infection. After infection, cells were washed 3 times with PBS and were incubated with overlay medium without suramin. After 3 days, cells were fixed, stained and plaques were counted. Suramin caused a dose-dependent reduction in the number of plaques and even at the lowest suramin concentration (12.5 µM) titers were already reduced by almost one log (Fig. 3C). These results suggest that suramin inhibits SARS-CoV-2 entry.

### Suramin inhibits SARS-CoV-2 replication in a primary human epithelial airway cell infection model

Primary human airway epithelial cell cultures (HAE) mimic the morphological and physiological features of the human conducting airway, arguably being the most relevant *ex vivo* model for human coronavirus research (23-25). For that reason, we decided to also evaluate the antiviral effect of suramin in this model. HAE were differentiated by culture at the air-liquid interface to achieve mucociliary differentiation, and were infected for one hour with 30,000 PFU of SARS-CoV-2 (estimated MOI of 0.1 based on the number of cells present on an insert), followed by washing with PBS. At 12 and 24 h.p.i., the cultures were treated on the apical side with either 50 µl of 100 µM suramin or 50 µl PBS. The HAE apical side was washed with PBS for 10 minutes at 37°C, and this supernatant was harvested at 12, 24 and 48 h.p.i. to analyze the viral load by RT-qPCR. RNA was also isolated from cells to quantify the levels of intracellular viral RNA and the housekeeping gene PGK1. RT-qPCR analysis of extracellular viral RNA levels showed that after infection approximately 10^7^ copies/ml of viral RNA remained at 1 h.p.i.. The viral load in the supernatant did not increase significantly at 12 and 24 h.p.i. in untreated cells, while at 48h a more than 1 log increase in viral RNA copies was observed. This is indicative of (very modest) viral replication in PBS-treated cells. The cultures that were treated with suramin displayed no increase in viral load in the supernatant, but rather even a slight decrease in copy numbers, suggesting viral replication did not progress in treated cells. At 48 h.p.i. the supernatant of suramin-treated cells showed 2-log lower SARS-CoV-2 released genome copy numbers than PBS-treated control cells (Fig. 4A). The levels of intracellular viral RNA displayed the same trend, with a decrease in viral RNA in suramin-treated samples compared to an increase in viral RNA in PBS-treated samples (Fig. 4B). A 1-log difference, from 10^6^ to 10^5^ copies per transwell was observed at 48 h.p.i. between suramin-and PBS-treated cells (Fig. 4B). The levels of the housekeeping gene, PGK1 remained stable in all samples, suggesting the reduction in viral RNA copies was not due to cell death. Moreover, cell viability measured by MTS assay (Fig. 4C) and LDH assay (data not shown), suggested suramin treatment (compared to PBS treatment) had no measurable cytotoxic effect on HAE cells. To determine the effect of suramin on infectious progeny released by HAE cells, we performed a plaque assay with the harvested supernatant. At 24 h.p.i., a modest difference was observed between the infectious progeny released by PBS (3.3 × 10^3^ PFU/ml) and suramin-treated cells (4.4 × 10^2^ PFU/ml). At 48 h.p.i., the supernatant of PBS-treated cells contained over 10^4^ PFU/ml, while no infectious virus was found in suramin-treated samples (Limit of detection 100 PFU/ml). This suggests that suramin reduces the progression of infection in a HAE culture infection model.

**Figure 4.**
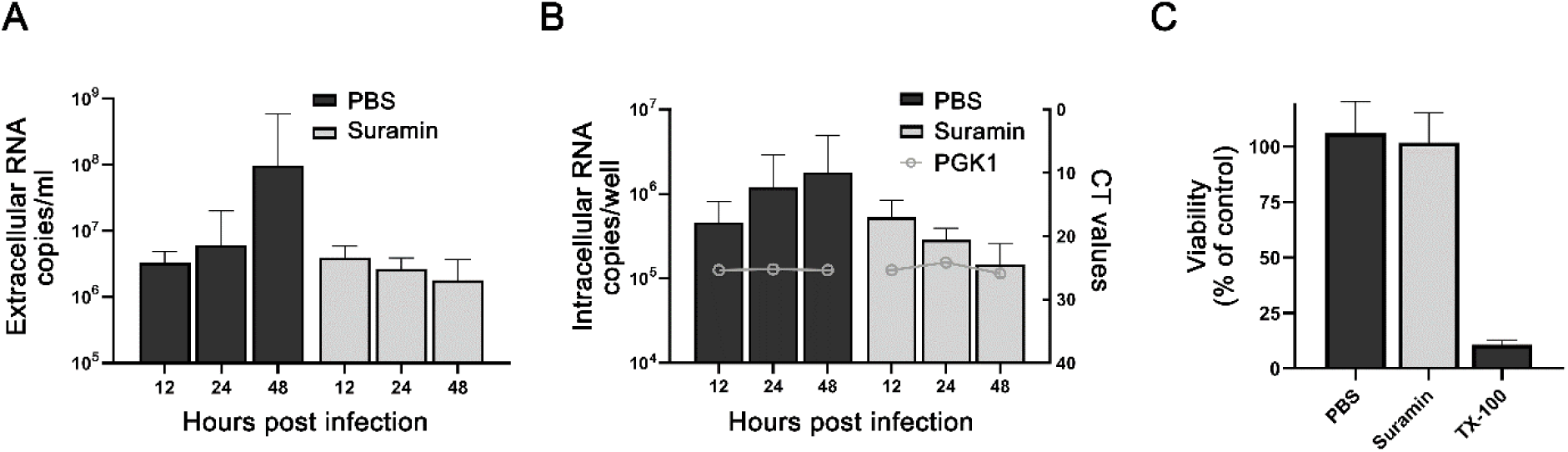
Suramin inhibits progression of SARS-CoV-2 infection in primary human airway epithelial cells. HAE cells were infected with 30,000 PFU of SARS-CoV-2 (estimated MOI of 0.1) and they were treated with 50 µl PBS or 50 µl of 100 µM suramin at 12 h.p.i. and 24 h.p.i.. (A) Levels of extracellular viral RNA were determined by RT-qPCR at 12, 24, and 48 h.p.i. (n=3). (B) Intracellular viral RNA levels were determined by RT-qPCR with an internally controlled multiplex (bars, left axis). Levels of the housekeeping gene PGK1 were analyzed to check for signs of cell death (gray lines, right axis). (C) Viability of suramin-treated cells evaluated by MTS assay, using treatment with 0.1% Triton X-100 as a positive control for cell toxicity (n=6).

## Discussion

The emergence of SARS-CoV-2 and its enormous impact on public health, society, economy and the lives of billions around the globe has prompted a multitude of efforts to develop vaccines and antivirals. Due to the lengthy development process of new and specific antivirals, there is a particular interest in repurposing existing drugs for treatment of the COVID-19 disease. This could provide a temporary solution, while better and more specific drugs are being developed. Several small-molecule compounds like chloroquine, hydroxychloroquine, favipiravir or remdesivir have been showing some efficacy against SARS-CoV-2 *in vitro* (26). However, despite promising results in preclinical studies, recent clinical trials (27, 28) suggested that these compounds, with the exception of remdesivir (8), do not provide much benefit to COVID-19 patients and could actually be dangerous due to possible side-effects. This leaves us currently empty-handed and in search for other approved drugs that might be repurposed. As an already approved antiparasitic drug, suramin would be one of the candidates for fast development of a treatment for COVID-19. Antiviral activity of suramin against RNA viruses was reported earlier by us and several other groups (15, 29-31) and the compound was and is also being evaluated in several clinical trials for other diseases, providing some evidence for its safety for therapeutic use. However, suramin can also cause several side-effects, which caused previous HIV trials with seriously ill patients to halt (32) and therefore caution is advised and it is crucial to conduct well controlled randomized trials, before any conclusions on possible benefits for COVID-19 patients can be drawn. Thus far, no studies have reported about a potential antiviral effect of suramin against coronaviruses.

In this study we assessed the antiviral activity of suramin against the newly emerged SARS-CoV-2. Suramin offered full protection against SARS-CoV-2-induced cell death in Vero E6 cells and inhibited the virus with an EC_50_ of 20 µM and a SI of >250 (Fig. 1). Suramin treatment of infected Vero E6 cells led to a reduction in extracellular viral RNA levels of up to 3 log. The highest concentration of compound that was used proved harmless to the cells and also previously cytotoxicity was only observed above 5 mM (15). Suramin also displayed antiviral efficacy in a human lung epithelial cell line and we observed a >2 log reduction in infectious virus progeny in suramin-treated cells.

Suramin was previously described to have the potential to inhibit several stages of virus replication by acting on different targets (15, 33). To assess which step in the SARS-CoV-2 replication cycle is affected by suramin treatment, we performed a time-of-addition assay. We observed that pre-treatment with suramin as well as addition during the first hour of infection resulted in a marked decrease of viral RNA in the supernatant, while treatments initiated after the first hour of infection showed no significant effect on virus replication, suggesting that suramin inhibits binding or entry. In addition, SARS-CoV-2 infectivity was decreased in plaque assays, when suramin was present only in the inoculum during infection, concordant with an effect on the early stages of infection (Fig. 3). This is in agreement with other studies that also reported on the inhibition of virus binding or entry by suramin (15, 30, 34). Our data suggest that the antiviral effect of suramin is primarily due to inhibition of binding and/or fusion.

Finally we evaluated the effect of suramin in a more relevant model of differentiated primary human airway epithelial (HAE) cells cultured and infected at the physiologically relevant air-liquid interface. We infected these cells with a relatively low dose of virus (estimated MOI of 0.1) and treated them with suramin by applying a 50 µl volume of 100 µM of suramin on the apical side at 12 and 24 h.p.i. This would allow us to follow spread of the viral infection and assess whether suramin is able to block progression of infection in this ‘treatment model’. HAE cell cultures are a composition of highly differentiated cells mainly containing basal, goblet, club and ciliated cells, hence representing an air-liquid interface that is mimicking the lung airway epithelium (35, 36). In a recent study, it was shown that SARS-CoV-2, like SARS-CoV, uses human angiotensin-converting enzyme 2 (ACE2) receptors for attachment in these human airway cells. Blocking of the host protease TMPRSS2, which is important for priming the fusion activity of the spike protein, also inhibited infection in lung cells (37). To address the variation of these proteins and the diversity of primary human airway cells within patients, we made use of HAE cultures that were obtained from different donors. Notably, we could observe differences in the susceptibility of cultures from different donors, in which HAE cultures from mixed donors showed higher titers. HAE cultures might have varying susceptibility to infection, possibly caused by a difference in cell differentiation and composition (38).

Administration of 100 µM of suramin on the apical side of the HAE cells did not appear to cause cytotoxic effects in our study (Fig. 4). In our HAE model for progression of SARS-CoV-2 infection, we infected cells at a low MOI and observed a modest (∼200-fold) increase in viral load by 48 h.p.i. in PBS-treated cultures. Although the increase in viral load was rather modest in control cells, we found no evidence for progression of the infection in suramin-treated cultures, as indicated by SARS-CoV-2 RNA levels that remained equal to that at 1 h.p.i. or even decreased over time. Moreover, the infectious progeny titer increased over time in PBS-treated HAE cultures and reached over 10^4^ PFU/ml by 48 h.p.i, while in suramin-treated HAE cells, infectious progeny showed a modest increase at 24 h.p.i. (10 fold lower than PBS-treated cells) and dropped to undetectable levels at 48 h.p.i. Since suramin-containing samples needed to be diluted by a 100-fold to exclude interference with the plaque assay, the limit of detection would be 100 pfu/ml. Even with this limit of detection, the supernatant collected from suramin-treated HAE cells contains at least 100 times less virus than that from PBS-treated cells. Much higher titers were obtained with HAE cultures from mixed donors than from single donors, but the inhibitory effect of suramin was also observed with single donor cultures. Overall, despite the modest level of infection in control cells, our results suggest that also in the HAE infection model, suramin has an inhibitory effect on progression of the SARS-CoV-2 infection.

Our study demonstrates that suramin inhibits SARS-CoV-2 replication in various cell culture models and at clinically achievable concentrations (after IV administration serum levels of >10x the EC_50_ could be achieved). Due to its mode of action (inhibition of entry) treatment of patients with suramin might require administration at an early stage, although it might also prevent spread of the virus in the lungs of already symptomatic patients or could prevent spread from respiratory tract to other organs. It might possibly even be used to prevent virus spreading in the nasopharynx, which appears to be the first site of infection (39-41). Standard treatment with suramin is done by intravenous administration, which would also be an option for seriously ill COVID-19 patients that are in intensive care, but is not ideal for other patients. As a negatively charged compound, suramin binds to various proteins and is poorly taken up by diffusion across the cell membrane, although it can be taken up by endocytosis (33). This poor uptake of suramin into cells might not necessarily be a problem for the efficacy against SARS-CoV-2, as it is expected to block the virus systemically and in the extracellular environment. Hypothetically, suramin administration into the respiratory tract in an aerosolized form could be even considered, although this requires new safety studies.

In conclusion, our preclinical study shows that suramin inhibits SARS-CoV-2 replication in cell culture, likely by preventing entry. Suramin also appears to prevent progression of SARS-CoV-2 infection in a human airway epithelial cell culture model. This is only the first step towards evaluating whether suramin treatment could provide any benefit to COVID-19 patients. Further studies should carefully evaluate different formulations, routes of administration, pharmacokinetics, and possible adverse effects in cell culture and *ex vivo* models. Ultimately, the clinical benefits of suramin for the treatment of COVID-19 patients should be evaluated in carefully performed and properly controlled clinical trials.

## Acknowledgements

The authors would like to thank Jessika Zevenhoven and Linda Boomaars-van der Zanden for technical assistance and Anne van der Does for helpful discussions. Clarisse S. B. da Silva was supported by the Coordination for the Improvement of Higher Education Personnel (CAPES) (Process nr. 88881.171440/2018-01), Ministry of Education, Brazil. Ying Wang was supported in part by a grant from the China Scholarship Council. Part of this research was supported by the Leiden University Fund (LUF), the Bontius Foundation and donations from the crowdfunding initiative “wake up to corona”.

## References

1. Lu H, Stratton CW, Tang YW. 2020. Outbreak of pneumonia of unknown etiology in Wuhan, China: The mystery and the miracle. J Med Virol 92:401–402.

2. Zhou P, Yang XL, Wang XG, Hu B, Zhang L, Zhang W, Si HR, Zhu Y, Li B, Huang CL, Chen HD, Chen J, Luo Y, Guo H, Jiang RD, Liu MQ, Chen Y, Shen XR, Wang X, Zheng XS, Zhao K, Chen QJ, Deng F, Liu LL, Yan B, Zhan FX, Wang YY, Xiao GF, Shi ZL. 2020. A pneumonia outbreak associated with a new coronavirus of probable bat origin. Nature 579:270–273.

3. Gorbalenya AE, Baker SC, Baric RS, de Groot RJ, Drosten C, Gulyaeva AA, Haagmans BL, Lauber C, Leontovich AM, Neuman BW, Penzar D, Perlman S, Poon LLM, Samborskiy DV, Sidorov IA, Sola I, Ziebuhr J, Coronaviridae Study Group of the International Committee on Taxonomy of V. 2020. The species Severe acute respiratory syndrome-related coronavirus: classifying 2019-nCoV and naming it SARS-CoV-2. Nature Microbiology 5:536–544.

4. WHO. 2020. World Health Organization, Coronavirus disease 2019 (COVID-19) Situation Report 106.

5. Weiss SR, Navas-Martin S. 2005. Coronavirus pathogenesis and the emerging pathogen severe acute respiratory syndrome coronavirus. Microbiol Mol Biol Rev 69:635–64.

6. Huang C, Wang Y, Li X, Ren L, Zhao J, Hu Y, Zhang L, Fan G, Xu J, Gu X, Cheng Z, Yu T, Xia J, Wei Y, Wu W, Xie X, Yin W, Li H, Liu M, Xiao Y, Gao H, Guo L, Xie J, Wang G, Jiang R, Gao Z, Jin Q, Wang J, Cao B. 2020. Clinical features of patients infected with 2019 novel coronavirus in Wuhan, China. The Lancet 395:497–506.

7. Wang M, Cao R, Zhang L, Yang X, Liu J, Xu M, Shi Z, Hu Z, Zhong W, Xiao G. 2020. Remdesivir and chloroquine effectively inhibit the recently emerged novel coronavirus (2019-nCoV) in vitro. Cell Res 30:269–271.

8. NIAID. 2020. National Institute of Allergy and Infectious Diseases. Adaptive COVID-19 Treatment Trial (ACTT). ClinicalTrialsgov Identifier: NCT04280705.

9. Voogd TE, Vansterkenburg EL, Wilting J, Janssen LH. 1993. Recent research on the biological activity of suramin. Pharmacol Rev 45:177–203.

10. Yahi N, Sabatier JM, Nickel P, Mabrouk K, Gonzalez-Scarano F, Fantini J. 1994. Suramin inhibits binding of the V3 region of HIV-1 envelope glycoprotein gp120 to galactosylceramide, the receptor for HIV-1 gp120 on human colon epithelial cells. J Biol Chem 269:24349–53.

11. Garson JA, Lubach D, Passas J, Whitby K, Grant PR. 1999. Suramin blocks hepatitis C binding to human hepatoma cells in vitro. J Med Virol 57:238–42.

12. Aguilar JS, Rice M, Wagner EK. 1999. The polysulfonated compound suramin blocks adsorption and lateral difusion of herpes simplex virus type-1 in vero cells. Virology 258:141–51.

13. Albulescu IC, Kovacikova K, Tas A, Snijder EJ, van Hemert MJ. 2017. Suramin inhibits Zika virus replication by interfering with virus attachment and release of infectious particles. Antiviral Res 143:230–236.

14. Basavannacharya C, Vasudevan SG. 2014. Suramin inhibits helicase activity of NS3 protein of dengue virus in a fluorescence-based high throughput assay format. Biochem Biophys Res Commun 453:539–44.

15. Albulescu IC, van Hoolwerff M, Wolters LA, Bottaro E, Nastruzzi C, Yang SC, Tsay SC, Hwu JR, Snijder EJ, van Hemert MJ. 2015. Suramin inhibits chikungunya virus replication through multiple mechanisms. Antiviral Res 121:39–46.

16. Yoshikawa T, Hill TE, Yoshikawa N, Popov VL, Galindo CL, Garner HR, Peters CJ, Tseng C-TK. 2010. Dynamic innate immune responses of human bronchial epithelial cells to severe acute respiratory syndrome-associated coronavirus infection. PloS one 5:e8729–e8729.

17. Kovacikova K, Morren BM, Tas A, Albulescu IC, van Rijswijk R, Jarhad DB, Shin YS, Jang MH, Kim G, Lee HW, Jeong LS, Snijder EJ, van Hemert MJ. 2020. 6′-β-Fluoro-homoaristeromycin and 6′-fluoro-homoneplanocin A are potent inhibitors of chikungunya virus replication through their direct effect on the viral non-structural protein 1. doi:10.1128/AAC.02532-19 %J Antimicrobial Agents and Chemotherapy:AAC.02532-19.

18. Schrumpf JA, Ninaber DK, van der Does AM, Hiemstra PS. 2020. TGF-β1 Impairs Vitamin D-Induced and Constitutive Airway Epithelial Host Defense Mechanisms. Journal of innate immunity 12:74–89.

19. Amatngalim GD, Schrumpf JA, Dishchekenian F, Mertens TCJ, Ninaber DK, van der Linden AC, Pilette C, Taube C, Hiemstra PS, van der Does AM. 2018. Aberrant epithelial differentiation by cigarette smoke dysregulates respiratory host defence. 51:1701009.

20. Kovacikova K, Morren BM, Tas A, Albulescu IC, van Rijswijk R, Jarhad DB, Shin YS, Jang MH, Kim G, Lee HW, Jeong LS, Snijder EJ, van Hemert MJ. 2020. 6′-β-Fluoro-Homoaristeromycin and 6′-Fluoro-Homoneplanocin A Are Potent Inhibitors of Chikungunya Virus Replication through Their Direct Effect on Viral Nonstructural Protein 1. 64:e02532–19.

21. Corman VM, Landt O, Kaiser M, Molenkamp R, Meijer A, Chu DKW, Bleicker T, Brünink S, Schneider J, Schmidt ML, Mulders DGJC, Haagmans BL, van der Veer B, van den Brink S, Wijsman L, Goderski G, Romette J-L, Ellis J, Zambon M, Peiris M, Goossens H, Reusken C, Koopmans MPG, Drosten C. 2020. Detection of 2019 novel coronavirus (2019-nCoV) by real-time RT-PCR. Euro surveillance: bulletin Europeen sur les maladies transmissibles = European communicable disease bulletin 25:2000045.

22. Scholte FEM, Tas A, Martina BEE, Cordioli P, Narayanan K, Makino S, Snijder EJ, van Hemert MJ. 2013. Characterization of Synthetic Chikungunya Viruses Based on the Consensus Sequence of Recent E1-226V Isolates. PLOS ONE 8:e71047.

23. Sheahan TP, Sims AC, Graham RL, Menachery VD, Gralinski LE, Case JB, Leist SR, Pyrc K, Feng JY, Trantcheva I, Bannister R, Park Y, Babusis D, Clarke MO, Mackman RL, Spahn JE, Palmiotti CA, Siegel D, Ray AS, Cihlar T, Jordan R, Denison MR, Baric RS. 2017. Broad-spectrum antiviral GS-5734 inhibits both epidemic and zoonotic coronaviruses. Sci Transl Med 9.

24. Sheahan TP, Sims AC, Zhou S, Graham RL, Pruijssers AJ, Agostini ML, Leist SR, Schafer A, Dinnon KH, 3rd, Stevens LJ, Chappell JD, Lu X, Hughes TM, George AS, Hill CS, Montgomery SA, Brown AJ, Bluemling GR, Natchus MG, Saindane M, Kolykhalov AA, Painter G, Harcourt J, Tamin A, Thornburg NJ, Swanstrom R, Denison MR, Baric RS. 2020. An orally bioavailable broad-spectrum antiviral inhibits SARS-CoV-2 in human airway epithelial cell cultures and multiple coronaviruses in mice. Sci Transl Med doi:10.1126/scitranslmed.abb5883.

25. Sims AC, Baric RS, Yount B, Burkett SE, Collins PL, Pickles RJ. 2005. Severe acute respiratory syndrome coronavirus infection of human ciliated airway epithelia: role of ciliated cells in viral spread in the conducting airways of the lungs. J Virol 79:15511–24.

26. Wang M, Cao R, Zhang L, Yang X, Liu J, Xu M, Shi Z, Hu Z, Zhong W, Xiao G. 2020. Remdesivir and chloroquine effectively inhibit the recently emerged novel coronavirus (2019-nCoV) in vitro. Cell Research 30:269–271.

27. Cao B, Wang Y, Wen D, Liu W, Wang J, Fan G, Ruan L, Song B, Cai Y, Wei M, Li X, Xia J, Chen N, Xiang J, Yu T, Bai T, Xie X, Zhang L, Li C, Yuan Y, Chen H, Li H, Huang H, Tu S, Gong F, Liu Y, Wei Y, Dong C, Zhou F, Gu X, Xu J, Liu Z, Zhang Y, Li H, Shang L, Wang K, Li K, Zhou X, Dong X, Qu Z, Lu S, Hu X, Ruan S, Luo S, Wu J, Peng L, Cheng F, Pan L, Zou J, Jia C, et al. 2020. A Trial of Lopinavir–Ritonavir in Adults Hospitalized with Severe Covid-19. doi:10.1056/NEJMoa2001282.

28. Molina JM, Delaugerre C, Le Goff J, Mela-Lima B, Ponscarme D, Goldwirt L, de Castro N. 2020. No evidence of rapid antiviral clearance or clinical benefit with the combination of hydroxychloroquine and azithromycin in patients with severe COVID-19 infection. Médecine et Maladies Infectieuses doi: https://doi.org/10.1016/j.medmal.2020.03.006.

29. Tan CW, Sam IC, Chong WL, Lee VS, Chan YF. 2017. Polysulfonate suramin inhibits Zika virus infection. Antiviral Research 143:186–194.

30. Chen Y, Maguire T, Hileman RE, Fromm JR, Esko JD, Linhardt RJ, Marks RM. 1997. Dengue virus infectivity depends on envelope protein binding to target cell heparan sulfate. Nature Medicine 3:866–871.

31. De Clercq E. 1979. Suramin: A potent inhibitor of the reverse transcriptase of RNA tumor viruses. Cancer Letters 8:9–22.

32. Kaplan LD, Wolfe PR, Volberding PA, Feorino P, Abrams DI, Levy JA, Wong R, Kaufman L, Gottlieb MS. 1987. Lack of response to suramin in patients with AIDS and AIDS-related complex. The American Journal of Medicine 82:615–620.

33. Wiedemar N, Hauser DA, Mäser P. 2020. 100 Years of Suramin. 64:e01168–19.

34. Ho Y-J, Wang Y-M, Lu J-w, Wu T-Y, Lin L-I, Kuo S-C, Lin C-C. 2015. Suramin Inhibits Chikungunya Virus Entry and Transmission. PLOS ONE 10:e0133511.

35. Hiemstra PS, Tetley TD, Janes SM. 2019. Airway and alveolar epithelial cells in culture. 54:1900742.

36. Fulcher ML, Randell SH. 2013. Human Nasal and Tracheo-Bronchial Respiratory Epithelial Cell Culture, p 109-121. *In* Randell SH, Fulcher ML (ed), Epithelial Cell Culture Protocols: Second Edition doi:10.1007/978-1-62703-125-7_8. Humana Press, Totowa, NJ.

37. Hoffmann M, Kleine-Weber H, Schroeder S, Krüger N, Herrler T, Erichsen S, Schiergens TS, Herrler G, Wu N-H, Nitsche A, Müller MA, Drosten C, Pöhlmann S. 2020. SARS-CoV-2 Cell Entry Depends on ACE2 and TMPRSS2 and Is Blocked by a Clinically Proven Protease Inhibitor. Cell 181:271-280.e8.

38. Menachery VD, Yount BL, Debbink K, Agnihothram S, Gralinski LE, Plante JA, Graham RL, Scobey T, Ge X-Y, Donaldson EF, Randell SH, Lanzavecchia A, Marasco WA, Shi Z-L, Baric RS. 2015. A SARS-like cluster of circulating bat coronaviruses shows potential for human emergence. Nature Medicine 21:1508–1513.

39. Lescure F-X, Bouadma L, Nguyen D, Parisey M, Wicky P-H, Behillil S, Gaymard A, Bouscambert-Duchamp M, Donati F, Le Hingrat Q, Enouf V, Houhou-Fidouh N, Valette M, Mailles A, Lucet J-C, Mentre F, Duval X, Descamps D, Malvy D, Timsit J-F, Lina B, van-der-Werf S, Yazdanpanah Y. 2020. Clinical and virological data of the first cases of COVID-19 in Europe: a case series. The Lancet Infectious Diseases doi: https://doi.org/10.1016/S1473-3099()30200-0.

40. Rockx B, Kuiken T, Herfst S, Bestebroer T, Lamers MM, Oude Munnink BB, de Meulder D, van Amerongen G, van den Brand J, Okba NMA, Schipper D, van Run P, Leijten L, Sikkema R, Verschoor E, Verstrepen B, Bogers W, Langermans J, Drosten C, Fentener van Vlissingen M, Fouchier R, de Swart R, Koopmans M, Haagmans BL. 2020. Comparative pathogenesis of COVID-19, MERS, and SARS in a nonhuman primate model. Science (New York, NY) doi:10.1126/science.abb7314:eabb7314.

41. Wölfel R, Corman VM, Guggemos W, Seilmaier M, Zange S, Müller MA, Niemeyer D, Jones TC, Vollmar P, Rothe C, Hoelscher M, Bleicker T, Brünink S, Schneider J, Ehmann R, Zwirglmaier K, Drosten C, Wendtner C. 2020. Virological assessment of hospitalized patients with COVID-2019. Nature doi:10.1038/s41586-020-2196-x.

